# High-throughput fitness profiling of Zika virus E protein reveals different roles for N-linked glycosylation during infection of mammalian and mosquito cells

**DOI:** 10.1101/199935

**Authors:** Danyang Gong, Tian-hao Zhang, Dawei Zhao, Yushen Du, Travis J. Chapa, Yuan Shi, Laurie Wang, Deisy Contreras, Gang Zeng, Pei-yong Shi, Ting-Ting Wu, Vaithilingaraja Arumugaswami, Ren Sun

## Abstract

Zika virus (ZIKV) infection causes Guillain-Barré syndrome and severe birth defects. ZIKV envelope (E) protein is the major viral protein involved in cell receptor binding and entry and therefore considered one of the major determinants in ZIKV pathogenesis. Here, we report a gene-wide mapping of functional residues of ZIKV E protein using a mutant library with changes covering every nucleotide position. By comparing the replication fitness of every viral mutant between mosquito and human cells, we identified that mutations affecting N-linked glycosylation at N154 position display the most divergence. Through characterizing individual mutants, we show that, while ablation of N-linked glycosylation selectively benefits ZIKV infection of mosquito cells by enhancing cell entry, it either had little impact on ZIKV infection on certain human cells or decreased infection through entry factor DC-SIGN. In conclusion, we define the roles of individual residues of ZIKV envelope protein, which contribute to ZIKV replication fitness in human and mosquito cells.

**Highlights:**

1. Gene-wide mapping of functional residues of E protein in human and mosquito cells.
2. Mutations affecting N-linked glycosylation display the most dramatic difference.
3. N-linked glycosylation decreases ZIKV entry into mosquito cells.
4. N-linked glycosylation is important for DC-SIGN mediated infection of human cells.

## Introduction

Zika virus (ZIKV) is a mosquito-borne human pathogen in the *Flaviviridae* family, which also includes the dengue (DENV), West Nile (WNV), Japanese encephalitis (JEV), and yellow fever viruses (YFV) (Brett D. Lindenbach, 2007). ZIKV was first isolated from the serum of a sentinel rhesus monkey in the Zika forest of Uganda in 1947 and was subsequently recovered from the mosquito *Aedes africanus* in the same forest (Dick et al., 1952). ZIKV infection is mostly asymptomatic, but can cause influenza-like symptoms such as fever, headache, joint pain and maculopapular rash (Duffy et al., 2009; Simpson, 1964). The recent outbreak of ZIKV in the Americas has demonstrated the potential for ZIKV to cause more serious disease, including microcephaly, other congenital malformations, and Guillain-Barré syndrome.

The ZIKV genome consists of a 10.8 kilobase single-stranded positive-sense RNA that codes for three structural proteins [capsid (C), membrane (prM/M), and envelope (E)] and seven non-structural proteins (NS1, NS2A, NS2B, NS3, NS4A, NS4B, and NS5). In addition, there are short untranslated regions (UTRs) on both the 5’- and 3’-ends of the genome (Kuno and Chang, 2007). The mature ZIKV virion is roughly spherical and approximately 50 nm in diameter. It contains a nucleocapsid that is surrounded by an icosahedral shell consisting of 180 copies of both E glycoprotein and M protein anchored in a lipid bilayer (Kostyuchenko et al., 2016; Sirohi et al., 2016). The flavivirus E protein, arranged as dimers on the surface of the mature virion, is the major viral protein involved in host-cell entry factor binding and fusion (Brett D. Lindenbach, 2007). Each E protein monomer consists of four domains—three ectodomains (DI, DII and DIII), and a transmembrane domain (TM). The structurally central DI acts as a bridge between DII and DIII, and contains one N-linked glycosylation site (N154). The N-linked glycosylation at residue 153/154 of the E protein is conserved across most flaviviruses, and has been shown to be important for optimal infection of mosquito and mammalian cells (Heinz and Allison, 2003; Lee et al., 2010; Post et al., 1992; Roehrig et al., 2007). DII includes the dimerization interface and a fusion loop that interacts with the endosomal membrane after conformation change. The IgG-like DIII is a continuous polypeptide segment and is thought to be important for binding to entry factors. Several host entry factors, including DC-SIGN, AXL, and TYRO3, have been shown to be important for mediating ZIKV infection (Hamel et al., 2015; Nowakowski et al., 2016). However, the detailed mechanism by which the E protein interacts with host-cell entry factors or the sequence determinants that contribute to human versus mosquito cell tropisms, is not fully known.

We have developed a high-throughput fitness profiling approach that combines high-density mutagenesis with the power of next-generation sequencing to identify functional residues in the context of virus infection (Qi et al., 2017; Qi et al., 2014; Remenyi et al., 2014; Wu et al., 2016). In this study, we applied this approach to systematically analyze the functional residues of ZIKV E protein during infection of mosquito and human cells. We achieved high sensitivity in identifying residues essential for ZIKV E protein function. Surprisingly, we found that N-linked glycosylation at position N154 had differential effects on ZIKV infection between mosquito and human cells. While ablation of this glycosylation had little impact on viral infection of human cells (A549 and hCMEC), it significantly increased infection of mosquito cells (C6/36), most probably by enhancing ZIKV entry. Lastly, N154 glycosylation was found to be important for ZIKV infection of mammalian cells through entry factor DC-SIGN, further broadening our current knowledge concerning the glycosylation of E protein in the mammalian and invertebrate ZIKV life cycles.

## Results

### Establishing infectious cDNA clone of ZIKV Strain PRVABC59

In order to facilitate mutational analysis of the ZIKV genome, we generated a plasmid (pZ-PR) carrying ZIKV cDNA which was generated from an early passage of PRVABC59 virus. The ZIKV PRVABC59 strain was isolated from a human serum specimen from Puerto Rico in December of 2015. The plasmid contains cDNA coding the entire ZIKV RNA genome, a T7 promoter on the 5’-end to drive ZIKV RNA *in vitro* transcription, and a BstBI site on the 3’-end to linearize the plasmid (Figs. 1A & 1B). For reconstituting ZIKV virions, 5’ capped viral genomic RNA was generated by *in vitro* transcription (Fig. 1C), and then electroporated into BHK21 cells. Culture supernatant containing recombinant ZIKV was harvested at 3 days post-electroporation, and further subjected to single-round amplification in C6/36 cells to generate the viral stock. Next, we compared recombinant and parental ZIKVs in cell culture, and found that there was no obvious difference in either the percentages of cells positive for ZIKV infection, or in the signal intensity of E proteins (Fig. 1D). In addition, they showed comparable growth kinetics in mammalian and mosquito cells (Fig. 1E). These results indicate that the recombinant ZIKV can be reconstituted from the ZIKV cDNA plasmid, and that it displays comparable infectivity and growth to the parental virus.

**Fig. 1.**
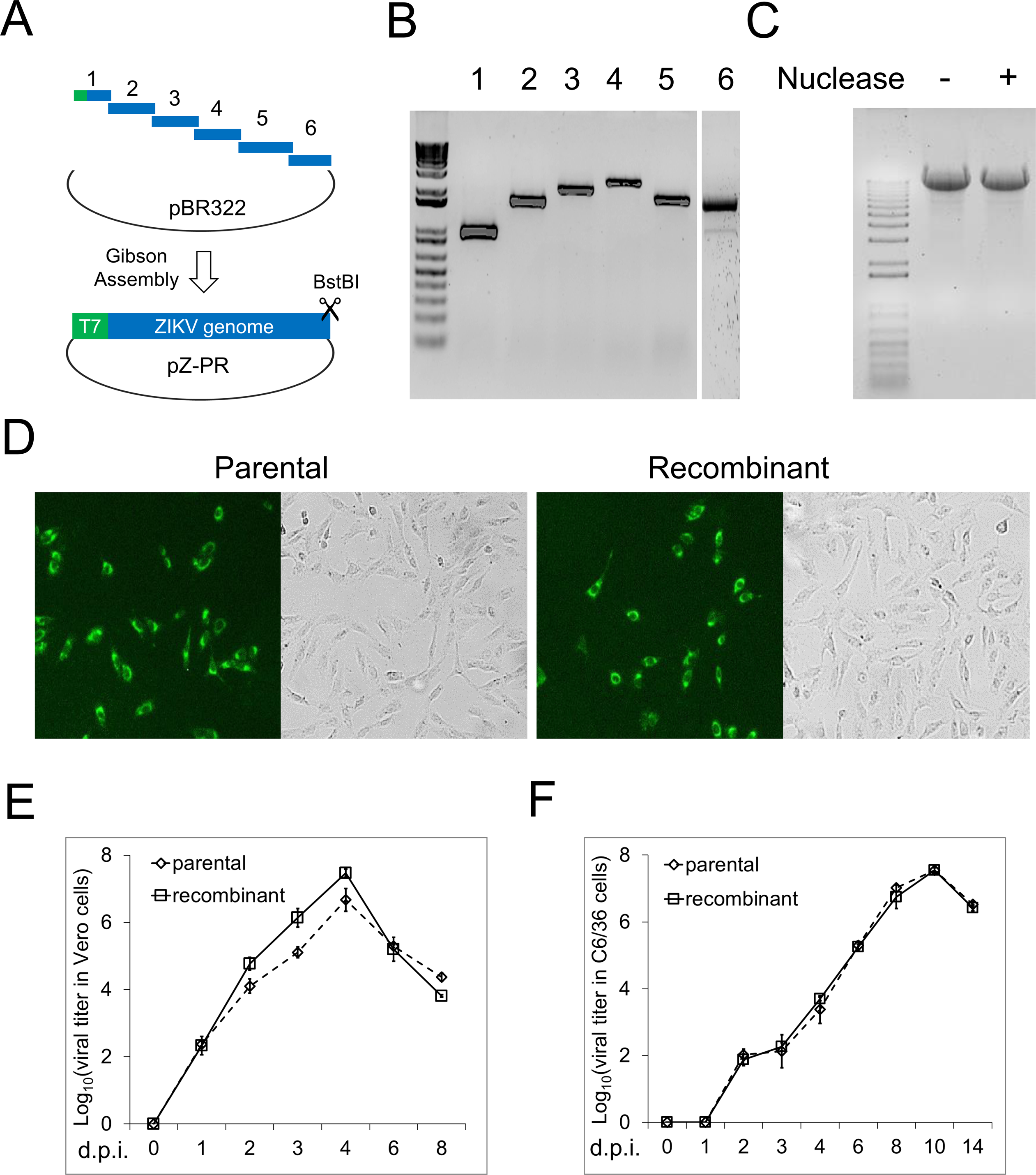
Construction of ZIKV infectious cDNA clone using strain PRVABC59. (A) Illustration of the cloning strategy. (B) Analysis of cDNA fragments on an agarose gel. The first 5 fragments were amplified from cDNA synthesized with random hexamer, and the 6^th^ fragment was generated from cDNA synthesized with specific oligo against the last 20 nt of ZIKV 3’-end. (C) Analysis of *in vitro* transcribed ZIKV RNA on a denaturing agarose gel. pZ-PR plasmids were linearized by BstBI, treated (or mock treated) with Mung Bean Nuclease to remove the two extra nucleotides, *in vitro* transcribed, and then separated on a 0.8% formaldehyde denaturing gel. (D) Immunostaining of ZIKV E proteins in parental or recombinant ZIKV infected cells. Vero cells were infected at a MOI of 0.5 and after 16 hrs, cells were fixed and subjected to IFA with αE (4G2) antibody. (E) Multiple-step growth curve analysis of recombinant and parental ZIKVs. Vero and C6/36 cells were infected at a MOI of 0.01, supernatants were harvested at the indicated time points post-infection to titrate virion production. Viral titers with error bars are plotted as mean ± SD (n=3).

### Constructing a library of mutations in ZIKV E protein

After generating the ZIKV cDNA clone, we aimed to systematically analyze the role of each residue of the E protein using a high-throughput fitness profiling approach. In this study, error-prone PCR was employed to introduce random point mutations into the ZIKV E protein coding region (1,512 nt) by using an enzyme blend to minimize the mutation bias (Fig. 2A). To better control the mutation rate of the error-prone PCR, the E protein was divided into three fragments, which together cover the whole sequence of the E protein. A mutant plasmid DNA library of ZIKV E protein was constructed by combining ∼40,000 clones from each fragment sub-library. To reconstitute the mutant virus library, the viral RNA library was generated from the plasmid DNA library by *in vitro* transcription, and then electroporated into BHK-21 cells.

**Fig. 2.**
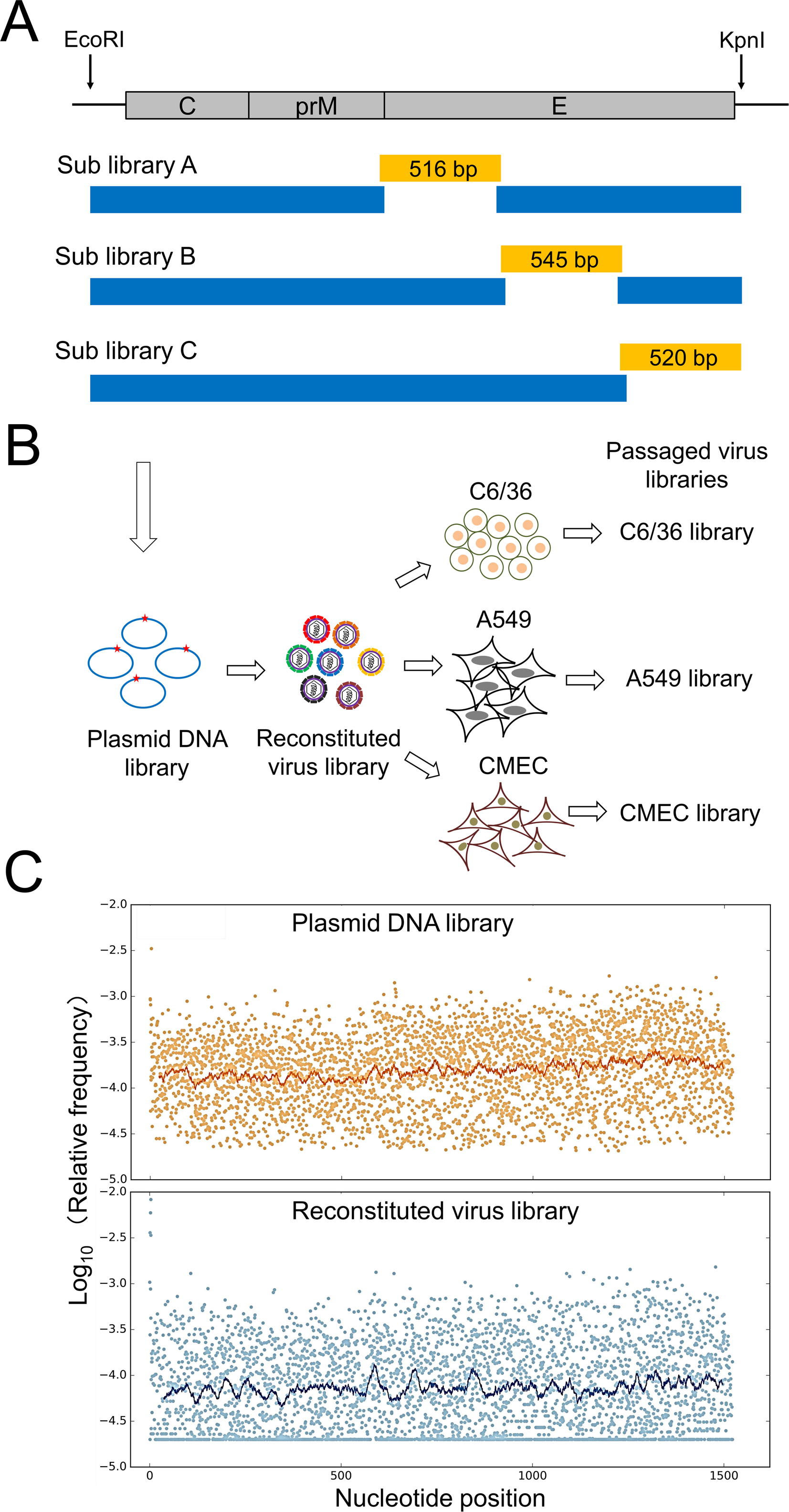
Construction and characterization of ZIKV E protein mutational library. (A) Construction of the plasmid DNA library. Three segments (yellow color) covering the entire E coding region were generated by error-prone PCR, while the corresponding left and right fragments (blue color) were generated by high fidelity PCR. Fragments were further assembled and cloned into pZ-PR plasmids. (B) A schematic representation of virus library construction and fitness profiling. Virus library was reconstituted by *in vitro* transcription from DNA library, followed by electroporation viral RNA into BHK21 cells. Virus library was then passaged in C6/36, A549 or hCMEC cells. Deep sequencing of plasmid DNA library, reconstituted virus library and passaged libraries was performed. (C) Relative frequencies of individual point mutations are plotted across the E protein. Mutations were examined by deep sequencing the ZIKV E gene, and the relative frequency of each mutation was calculated by dividing the mutation occurrence by the total population count (WT plus mutations). A smooth curve was fitted by cross correlation and plotted.

In order to evaluate the quality of the ZIKV E mutant libraries, both the plasmid DNA library and the reconstituted virus library were subjected to deep sequencing analysis. In addition, we sequenced a technical replicate for the DNA library, and triplicates for the virus library, to estimate the reproducibility of individual steps (Table S2). In the DNA library, 99.27% (4,503 out of 4,536) of expected mutations were identified by deep sequencing, in which 3,849 mutations showed average relative frequency higher than 20 per million reads (Table S3). Here, the relative frequency of each mutation was determined by dividing its mutation read count by the total population read count (see Material and Methods). These 3,849 mutations covered all 1,512-nt positions of the ZIKV E protein, and could be categorized as 2,729 missense (70.88%), 928 silent (24.11%), 192 nonsense (4.99%) mutations. The 2,729 missense mutations introduced 2,494 unique amino acid changes, and covered all residues of ZIKV E protein. In addition, mutations showed relatively even distributions across the E protein coding sequence (Fig. 2C) and no strong bias on the nucleotide composition (Table S4). As error-prone PCR enzyme blend introduces mutations randomly, the frequency of mutations in the plasmid DNA library should follow Poisson distribution, where all mutation events are independent and random. Based on the deep sequencing result from the DNA plasmid library, each individual viral genome has 0.82 mutations on average. 44.2% of the genomes are expected to be wild type, 36.1% of the genomes are expected to have a single mutation, and 14.7% of the genomes are expected to have two mutations. Only 5.0% of the genomes are expected to have more than two mutations. In the reconstituted virus library, 93.56% of expected mutations were identified, with 2,860 mutations showing average relative frequency greater than 20 per million reads. Compared to the plasmid DNA library, these mutations included similar numbers of silent (908 and 31.75%), slightly fewer missense (1,845 and 64.51%), and significantly fewer nonsense mutations (107 and 3.74%) (Table S3). The 1,845 missense mutations introduced 1,753 unique amino acid changes, and covered 503 out of the 504 residues of the ZIKV E protein (except Gly5). These results clearly show that we have generated a high-quality plasmid DNA library, and that we have reconstituted a virus library with high-complexity. To evaluate the reproducibility of experimental procedures, we examined the frequency distribution of all mutant types across replicate libraries, and we observed similar distribution between both of the DNA libraries and among the three virus libraries (Fig. S1A). In addition, we compared the relative frequency of individual mutations between replicate libraries. A Pearson’s correlation coefficient of 0.97 was obtained for the technical replicates of the DNA library, and 0.89 or 0.88 for the virus libraries (Fig. S1B). These strong correlations validated the reproducibility of our high-throughput fitness profiling platform.

To collect sufficient quantities of ZIKV mutant viruses for subsequent experiments, our virus library was harvested at four days after RNA electroporation. Trans-complementation after RNA transfection should enable the reconstitution of most mutant viruses. Due to some viral amplification, more likely in the late period, there could be some selection effect on mutants. To evaluate the potential selection effect on the virus library, we calculated the relative fitness score (RF score is expressed as Relative frequency_*virus*_/Relative frequency_*DNA*_) for each mutant in the virus library. As shown in Fig. S1C, the RF score of silent mutations (mean=0.88) was higher than those of nonsense (mean=0.06) and missense (mean=0.21) mutations, suggesting that some fitness selection did occur during the virus reconstitution step. However, the high-complexity of missense mutations in the virus library, and the depth of our sequencing, can support a high-resolution fitness profiling of ZIKV E protein in the three culture cell lines.

### Fitness profiling of E protein mutants in mosquito and human cells

The ZIKV E protein mediates viral entry into host cells, defines tropism, and is an attractive target to counteract infection. As an arbovirus, the life cycle of ZIKV involves transmission between *Aedes* mosquitoes and human hosts (Ayres, 2016; Marchette et al., 1969). Therefore, it is important to identify host-specific functional residues of the E protein, which in turn can facilitate studies of the viral life cycle, especially regarding its entry mechanisms. In this study, one mosquito (C6/36) and two human (A549 and hCMEC/D3) cell lines were employed for the following reasons: 1) C6/36 cells were generated from the larvae of *Aedes albopictus* and have been widely used to study ZIKV and other flaviviruses; 2) Epithelial cells are natural targets for ZIKV infection (Ma et al., 2016; Singh et al., 2017; Tang et al., 2016b), and as a human epithelial cell line, A549 expresses a majority of reported ZIKV entry factors (except DC-SIGN) (unpublished RNAseq result) supporting efficient infection of ZIKV; 3) ZIKV targets neuro cells by passing the human blood-brain barrier (Miner and Diamond, 2017; Nowakowski et al., 2016; Tang et al., 2016a), and barrier endothelial cells are permissive for ZIKV infection (Liu et al., 2016; Richard et al., 2017; Singh et al., 2017); thus, hCMEC/D3, which is a human blood-brain barrier endothelial cell line, was also chosen for our study. We first examined the fitness of E protein mutants by passaging the reconstituted virus library in these three cell lines respectively (Fig. 2B and Table S3), and noticed that the RF score distributions of silent mutations were clearly different from that of nonsense and missense mutations (Fig. S2), which indicated a fitness selection in all three cell lines. It is notable that there are some nonsense mutations identified in the passaged virus libraries. In order to have rapid adaptation and evolution, RNA viruses have an average mutation rate (closely equivalent to mutational frequency) of 10^-4^ -10^-5^ mutations per nucleotide (nt) per round of RNA replication (Domingo et al., 1996; Drake et al., 1998; Meyerhans, 1999; Ramirez, 1995). In our passaged virus libraries from C6/36, A549 and CMEC cells, the mean frequency of nonsense mutations is 3.08×10^-5^, 3.42×10^-5^ and 3.23×10^-5^ respectively, which is consistent with the spontaneous mutation rates that occur during genome replication. Therefore, we conclude that nonsense mutations identified in our virus libraries were most likely generated by spontaneous mutations instead of contamination by transfected RNA.

As a control for our fitness profile, we evaluated residues that have high variability in nature. We downloaded all ZIKV E protein sequences from Virus Pathogen Resource, and extracted 52 unique complete sequences. After calculating the polymorphism score of each residue, we found that nine residues were highly diverse with a SNP score ≥ 100 (Table S5 and Fig. 3A). The high diversity of these residues may suggest the ability of tolerating mutations. Indeed, the average RF scores of mutations on these nine residues was close to 1 in all three cell lines (Fig. 3B). Moreover, we found that the frequently occurring natural variants on these nine residues did not or only slightly caused fitness loss (Fig. 3C).

**Fig. 3.**
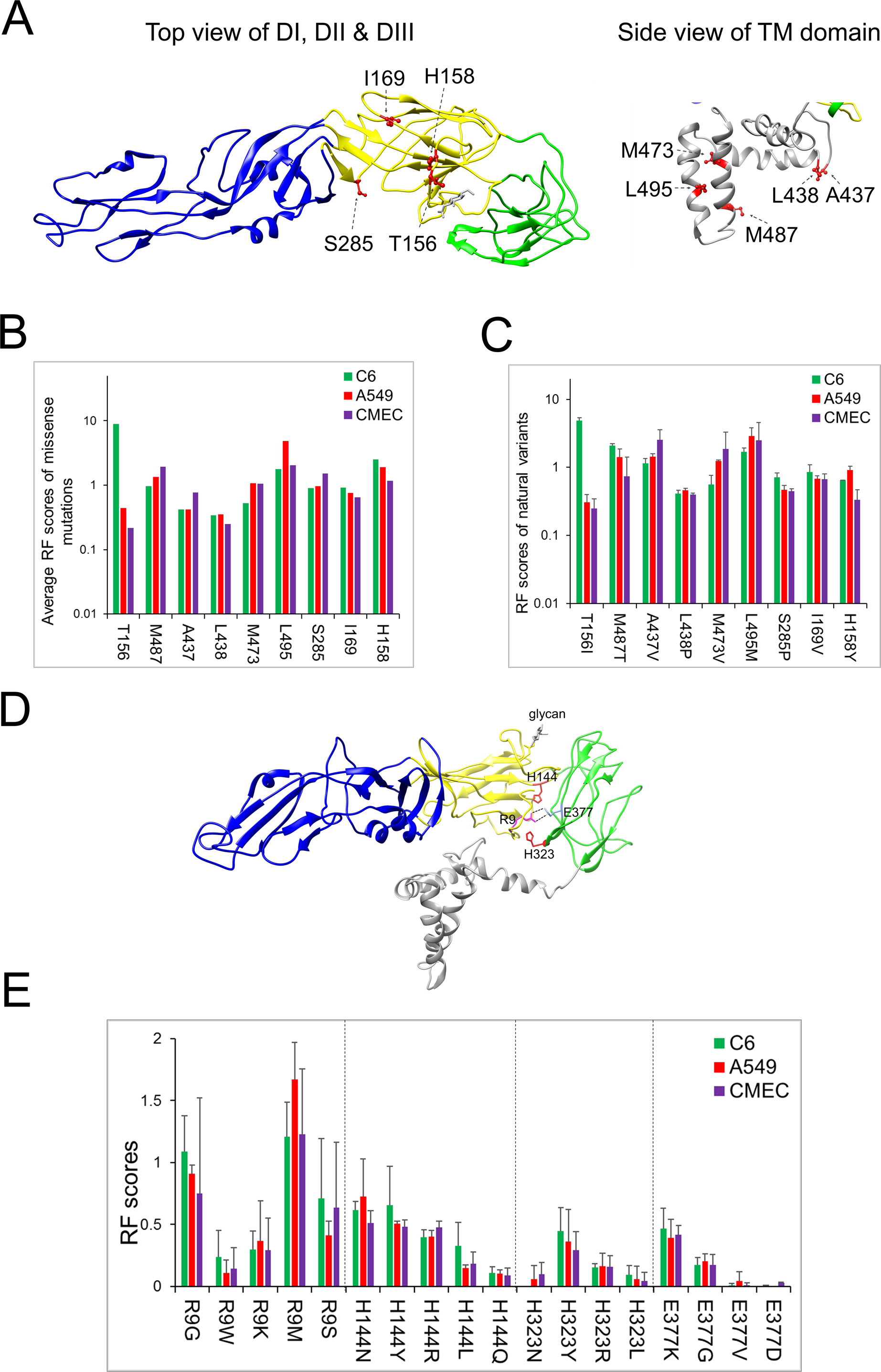
Analysis of residues highly polymorphic or involved in conserved function. (A) Ribbon diagrams of E protein monomer and localization of 9 residues highly polymorphic in natural ZIKV sequence. Yellow, domain I; Blue, domain II; Green, domain III; Grey, transmembrane domain. (B) Average relative fitness score (RF score) of mutations on residues. The RF score of mutation is expressed as the ratio of relative frequency in viral library to the relative frequency in DNA library. The average RF score of mutations on residue is defined as an average of RF scores of all missense mutations on this residue. (C) RF scores of natural occurring mutations. (D) Localization of 4 conserved residues potentially functioning in pH-induced conformational change. A salt bridge between R9 and E377 is highlighted. (E) RF scores of mutations on residues R9, H144, H323 and E377.

To guide our analysis of the fitness data, we focused on the pH-induced conformation change of E protein. Previous studies have shown that H146 and H323 in tick-borne encephalitis virus (equivalent to H144 and H323 in ZIKV) functioned as molecular switches (Fig. 3D) (Fritz et al., 2008; Prakash et al., 2010).Additionally, R9 in domain I (DI), that E373 (E377 in ZIKV) in domain III (DIII) form a salt bridge through their side chains in neutral pH, and that this interaction is essential for the dimeric conformation of E in mature virions (Fig. 3D). After extensive conformation change triggered by low pH, a different salt bridge forms between E373 and H323 which may contribute to the stability of the post-fusion trimer (Bressanelli et al., 2004). After analyzing the fitness of mutations on these 4 residues, we found that: 1) every mutation showed comparable fitness among the three cell lines; 2) all mutations on H323 and E377 showed RF scores lower than 0.5, and four specific mutations (H323N, H323L, E377V and E377D) reduced RF scores to <0.1; 3) in contrast, five mutations on R9 and H144 (R9G, R9M, R9S, H144N and H144Y) showed RF scores greater than 0.5, and no mutation on these two residues reduced RF scores below 0.1 (Fig. 3E). These results suggest that H323 and E377 are important for the survival of ZIKV independent of cell type, while R9 and H144 play relatively minor roles. In addition, the above findings validate the sensitivity of our high-throughput fitness profiling approach.

Next, we plotted the RF scores of mutations across the entire ZIKV E protein (Fig. 4A), and found that the majority of mutations resulted in comparable fitness among the original reconstituted virus and the three passaged libraries. Interestingly, there was a cluster of beneficial mutations in C6/36 cells around the N-linked glycosylation site (N154). Smoothed local average analysis further confirmed that mutations in the glycosylation region presented the most dramatic difference in comparable fitness, where they showed significantly higher fitness in mosquito C6/36 cells but not in the two human cells tested (Fig. 4B).

**Fig. 4.**
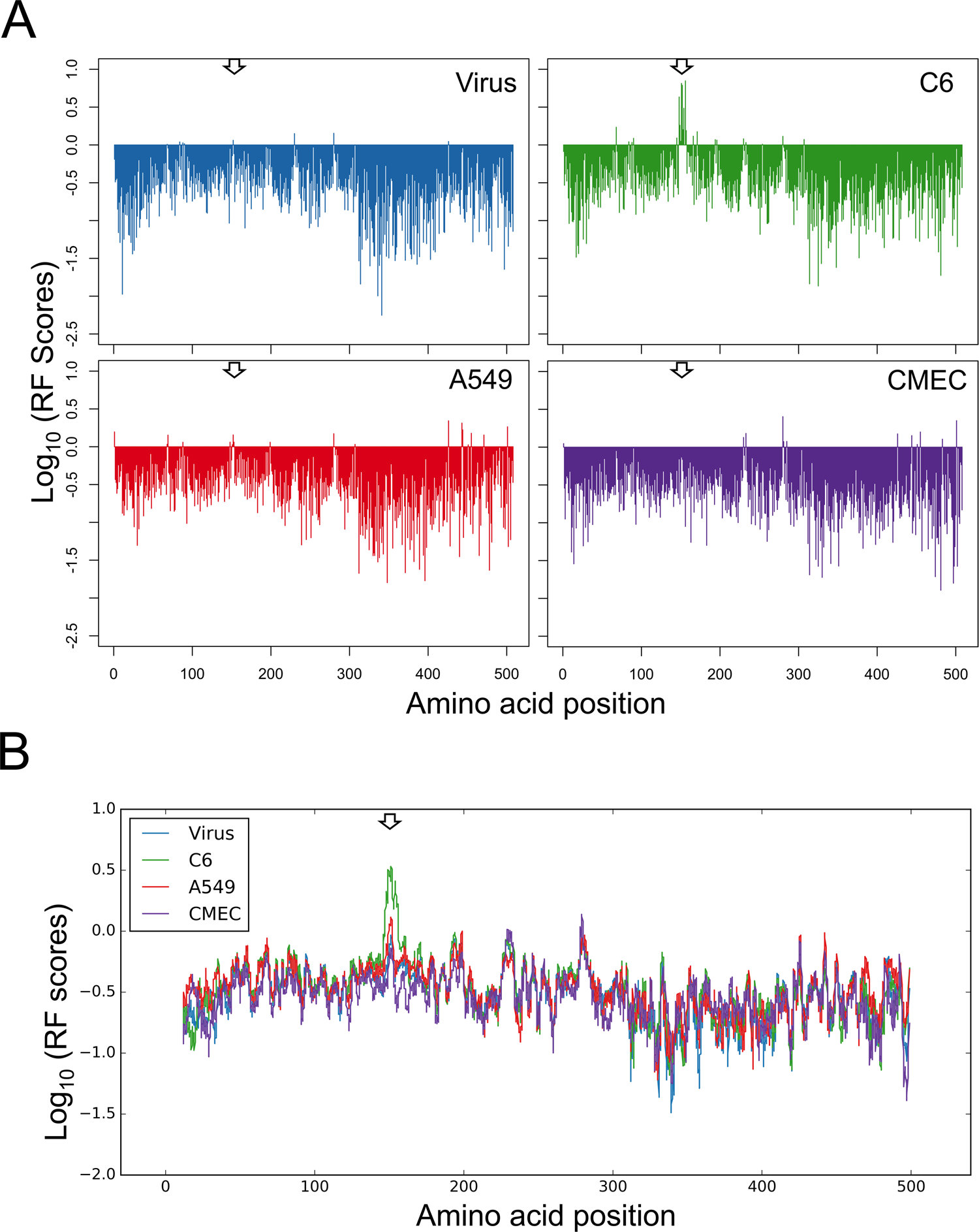
Fitness profiling of ZIKV E protein mutant virus libraries. (A) Plot of averaged relative fitness scores (RF scores) of mutations at each residue position. (B) Smoothed local average plot of mutants’ RF scores. Smoothed local average was expressed as the rolling mean of the mutations’ RF scores with window size of 20. Arrow, N-linked glycosylation site (N154).

### Constructing and characterizing individual N-linked glycosylation defective mutants

Previous studies on the cryoEM structure of ZIKV mature virion showed that the E protein was glycosylated at N154 position (Kostyuchenko et al., 2016; Sirohi et al., 2016). This is consistent with the knowledge that N-linked glycosylation requires a N-X-S/T consensus sequence (Taylor and Drickamer, 2011), which is present as N154-D155-T156 in ZIKV stain PRVABC59. In our deep sequencing analysis, many mutations on or around N154 position were beneficial for ZIKV replication in mosquito C6/36 cells, likely due to ablation of N-linked glycosylation. As shown in Fig. 5A, all silent mutations (I152I, V153V, N154N, D155D and T156T) did not obviously affect viral fitness, further validating our high-throughput screening approach. Eleven missense mutations were identified on two residues, N154 and T156, and in general they selectively increased ZIKV fitness in C6/36 cells by 10 folds. Additionally, four missense mutations on I152 (I152V, I152N and I152T) and V153 (V153D) caused a similar phenotype. Taken together, the profiling results suggest that glycosylation removal at N154 of ZIKV E protein might contribute to an increased viral fitness in mosquito C6/36 cells.

**Fig. 5.**
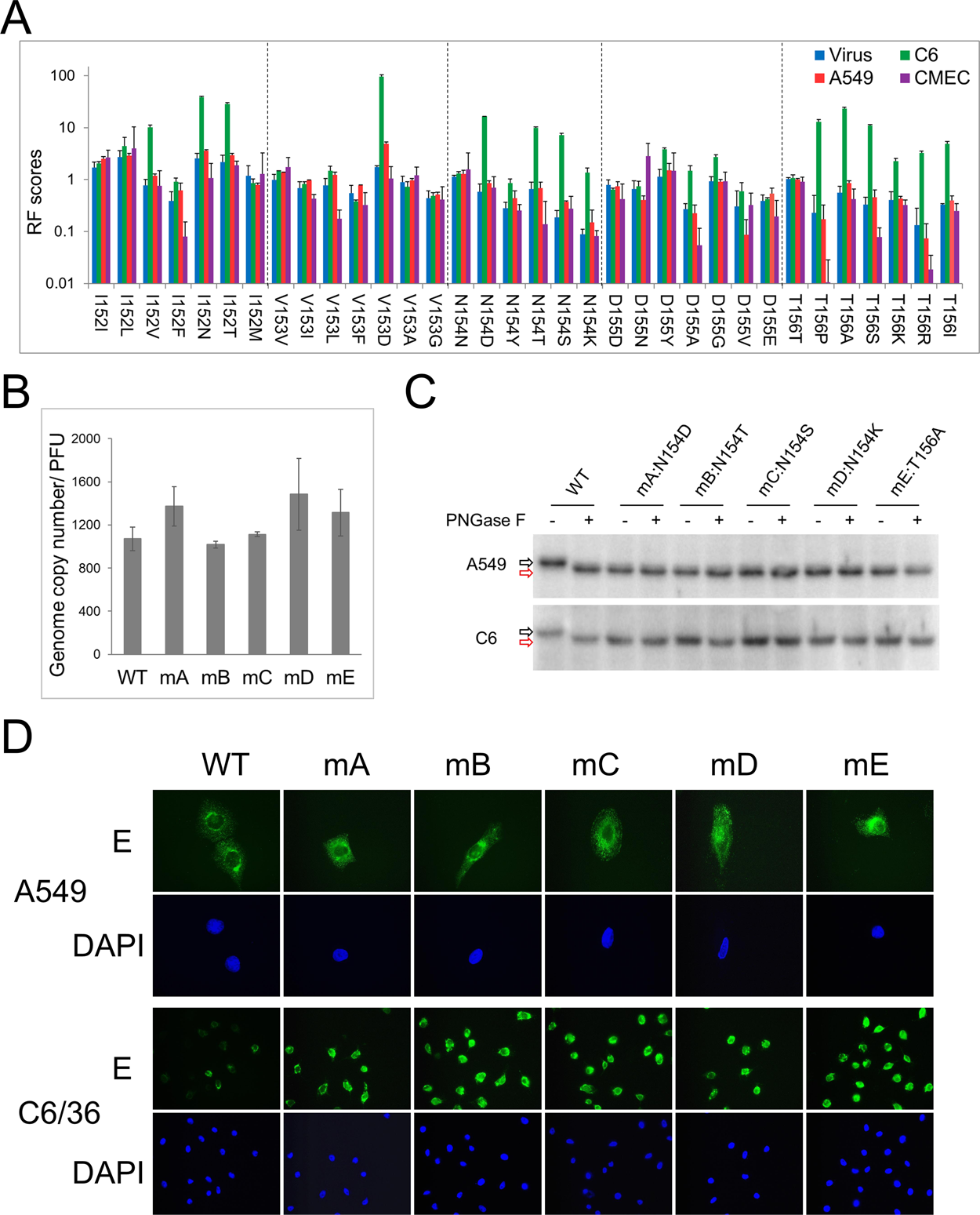
Analysis of mutations in the N-linked glycosylation region. (A) RF scores of mutations covering the five residues on and surrounding the N154 position. (B) Infectivity of ZIKV WT and mutants. Titers of ZIKV WT and mutants were determined by plaque forming assay on A549 cells, while genome copies of the same viruses were quantified by qRT-PCR. The ratio of viral genome copies to PFU was plotted as mean ± SD (n=3). (C) Analysis of E protein N-linked glycosylation using PNGase F treatment and western blotting. Black arrow, N-linked glycosylation; Red arrow, absence of N-linked glycosylation. (D) Cells from same infection as (C) were also subjected to immunostaining using αE (4G2) antibody.

To confirm the profiling results and to explore the mechanism of how N-linked glycosylation impacts ZIKV fitness, five point mutations, four on N154 and one on T156, were individually introduced into the pZ-PR plasmid (Table S1). ZIKV mutants were reconstituted in BHK21 cells, and were further amplified in C6/36 cells to generate viral stocks. In plaque forming assays on A549 cells, all mutants generated plaques of comparable size to wild type (WT) ZIKV (data not shown). To quantify the infectivity of ZIKV WT and mutants, we calculated the particle-to-PFU ratios by utilizing genome copy number and the results of plaque assay (Fig. 5B). A ratio of 1.0x10^3^ was obtained for the WT virus, and ratios between 1.0 x10^3^ and 1.4 x10^3^ were obtained for the mutants, suggesting comparable infectivity among WT and mutants.

To test whether these five mutations affected the N-linked glycosylation, we employed Peptide-N-Glycosidase F (PNGase F) treatment and western blotting. As shown in Fig. 5C, PNGase F treatment increased the mobility of WT E protein, suggesting the removal of N-linked glycosylation. All mutant E proteins displayed the same mobility as PNGase F treated WT E protein, and were not sensitive to PNGase F treatment, indicating that all five mutations abolished N-linked glycosylation. We further examined the E protein localization and expression by IFA, and found that E proteins from ZIKV WT and mutants showed similar localization and expression levels during infection of A549 cells (Figs. 5D & S3). However, when infecting C6/36 cells, ZIKV mutants expressed more E proteins than WT virus, which is consistent with the results from western blotting (Figs. 5C& S3).

### Ablation of N-linked glycosylation benefits ZIKV replication in mosquito cells by enhancing virus entry

To evaluate the roles of the N-linked glycosylation during ZIKV infection of mosquito and human cells, we characterized the growth kinetics of WT and mutant viruses by quantifying intracellular viral genome replication (Figs. 6A and S4A) and extracellular infectious virion production (Figs. 6B and S4B). The data showed that all viruses grew comparably in A549 cells. In contrast, the growth of mutant viruses was selectively enhanced in C6/36 cells. The genome copies of mutant viruses were ∼10 times higher than that of WT virus at days 1 and 2 post infection,leading to ∼10 times more virion release at days 2 and 3 post infection. These results clearly demonstrate that ablation of N-linked glycosylation specifically benefited ZIKV growth in C6/36 cells, but not in A549 cells, validating our findings from the high-throughput fitness profiling analysis. Furthermore, we found that mutant viruses also showed significantly enhanced growth (Fig. S5) in *Aedes aegypti* CCL-125 cells, indicating that this phenotype may be common to mosquito cell lines but not something specific to C6/36 cells. However, we also noticed that at late times post infection of C6/36 cells, WT and mutant viruses showed similar genome replication and virus production (day 4 in Figs. 6A & 6B, days 4 and 6 in Fig. S4). One possible reason for this observation may be that infected C6/36 cells do not undergo cell lysis; thus, once the monolayer is completely infected, fitness advantages become no longer apparent. At 4 and 6 days post infection, all C6/36 cells were positive for ZIKV infection, maintained similar levels of intracellular viral genomes, and released comparable amounts of virions. Therefore, a possible mechanism that explains why ZIKV mutants show higher fitness in C6/36 cells is that ablation of N-linked glycosylation might enhance the early events of virus infection, including attachment and entry.

**Fig. 6.**
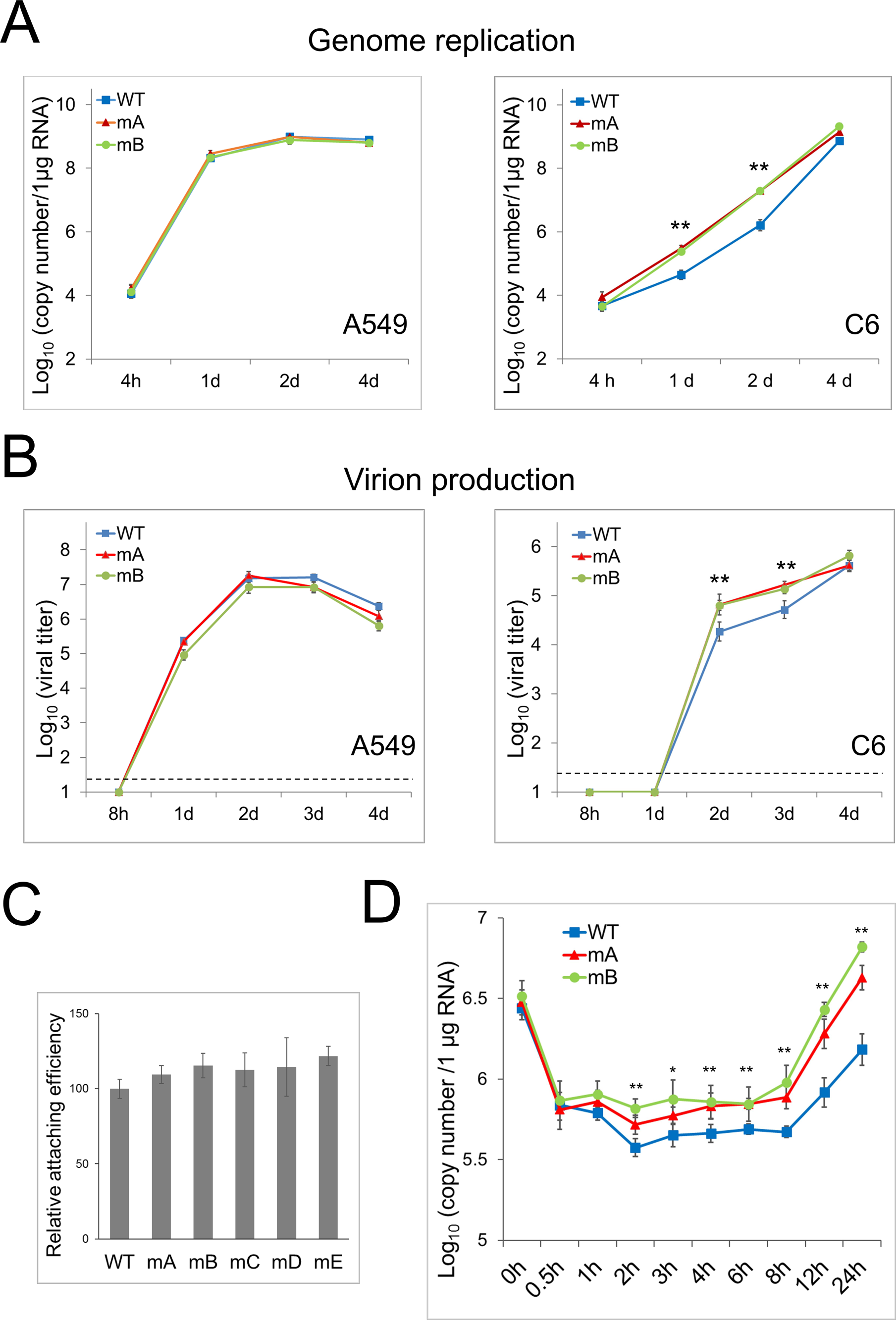
Characterization of ZIKV N-linked glycosylation defective mutants. (A) Intracellular genome replication of ZIKV WT and mutants. A549 or C6/36 cells were infected with ZIKV WT, mA or mB at a MOI of 0.05, harvested at indicated times for RNA purification and qRT-PCR analysis. (B) Infectious virus production. Supernatants were harvested from same cells as (A) to determine viral titers. (C) Attachment efficiency of ZIKV WT and mutants on surface of C6/36 cells. C6/36 were incubated with ZIKVs at a MOI of 10 for 1.5 h at 4 °C, followed by washing with ice cold PBS three times, then immediately subjected to RNA purification and qRT-PCR to determine viral genome copies. Relative attaching efficiency to WT virus was plotted as mean ± SD (n=3). (D) Entry efficiency of ZIKV WT and mutants. C6/36 cells were incubated with ZIKVs as (C). After washing with ice cold PBS, cells were replenished with warm culture medium before returning to 28 °C, harvested at indicated times, and subjected to qRT-PCR analysis.

To test this hypothesis, we first quantified the efficiency of ZIKV attachment and found that mutant viruses bound to the cell surface as efficiently as WT virus did (Figs. 6C & S6). Next, we examined the efficiency of viral genome delivery into the cytoplasm by quantifying cell-associated viral RNA. C6/36 cells were absorbed with WT or mutant viruses at 4 °C, and then returned to 28 °C to initiate virus entry followed by viral protein expression and genome replication. Cells were collected at various times post infection to quantify the viral genome. Right after 4 °C incubation and washing, cell-associated genome copies of WT and mutant viruses were similar (Fig. 6D, 0 h). At 0.5h after returning to 28 °C, despite an overall drop of ∼80%, the genome copies of all viruses were still comparable (Fig. 6D, 0.5h). It is likely that ZIKV loses a large portion of virions in acidic endosomes, as was reported for DENV (van der Schaar et al., 2007), which would explain the drop of viral genome copies between 0h and 0.5h. Genome copies of WT virus kept decreasing until 2h, and then maintained a steady level until 8h, a level which may reflect the amount of viral genome successfully delivered to cytoplasm. In contrast, genome copies of mutant viruses did not obviously change between 0.5h and 8h. Starting from 1h, mutant viruses consistently showed higher genome copies than WT virus, implying a higher efficiency of viral genome delivery. After 8h, genome copies of WT and mutant viruses robustly increased, indicating successful genome replication in each. Taken together, during ZIKV infection of mosquito cells, N-linked glycosylation seems to impair virus entry, resulting in less viral genome delivery into the cytoplasm than in non-glycosylated mutant viruses.

### N-linked glycosylation is important for ZIKV infection of mammalian cells through DC-SIGN

The conserved N-D-T sequence motif was identified in the majority of ZIKV strains (Faye et al., 2014; Lanciotti et al., 2008), suggesting that N-linked glycosylation of E protein should play some role(s) in certain step(s) of the ZIKV life cycle, though we did not identify using the above *in vitro* cell systems. DC-SIGN, also known as CD209, mediates DENV infection of dendritic cells by binding E protein glycan(s) (Navarro-Sanchez et al., 2003; Pokidysheva et al., 2006; Tassaneetrithep et al., 2003). Furthermore, expression of DC-SIGN makes cells susceptible to ZIKV infection (Hamel et al., 2015). Therefore, by using the mutant viruses, we sought to test whether the N-linked glycosylation of E protein was required for ZIKV infection of DC-SIGN expressing cells. Because 293T cells are not susceptible to ZIKV infection due to the lack of existing entry factor(s) (Hamel et al., 2015), we first established DC-SIGN expressing cells by transfecting 293T cells with DC-SIGN expression plasmid. As shown in Fig. S7, control 293T cells were largely non-permissive to ZIKV infection. The expression of DC-SIGN strongly enhanced WT ZIKV infection, resulting in ∼10% cells positive for ZIKV E protein. Further, we tested the infection efficiency of ZIKV WT and mutants by using the DC-SIGN expressing 293T cells. The results showed that when compared to WT virus, infection with mutant viruses resulted in significantly less E protein positive cells (Fig. 7A), as well as an ∼80% decrease of viral genome replication (Fig. 7B). Moreover, similar results were also observed during ZIKV infection of human immature dendritic cells (Figs. 7C & 7D). These observations suggest that N-linked glycosylation of E protein plays an important role in ZIKV infection through entry factor DC-SIGN.

**Fig. 7.**
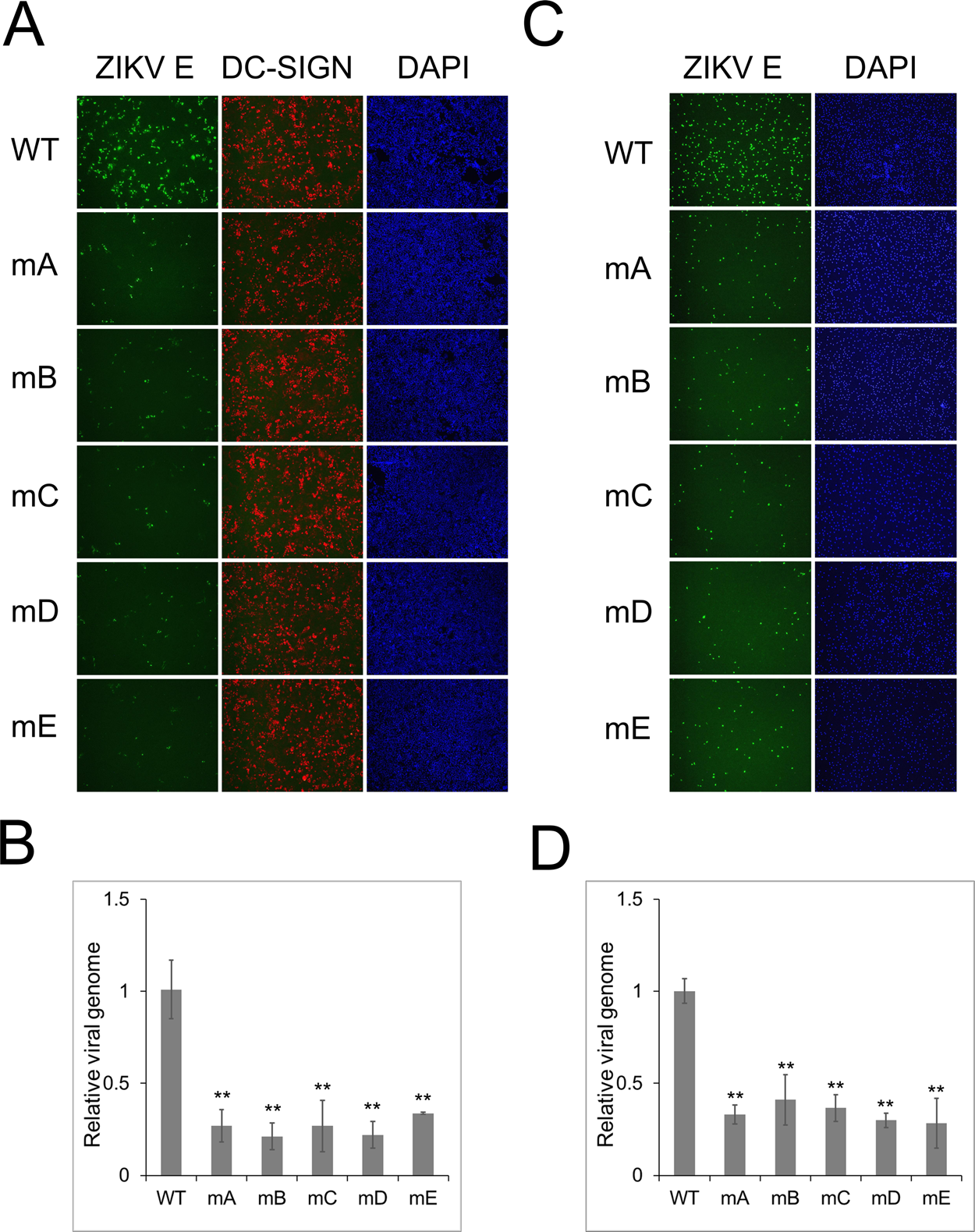
N-linked glycosylation enhances ZIKV infection through entry factor DC-SIGN. (A) Immunostaining of ZIKV E protein. DCSIGN-cFLAG transfected 293T cells were infected with ZIKV WT or mutants at a MOI of 0.2 for 16 hrs, then fixed and subjected to IFA analysis with αE (4G2) and αFLAG antibodies. (B) Total RNA was purified from same cells as (A), and subjected to qRT-PCR analysis to quantify viral genome copies. (C) Immunostaining of ZIKV E protein in immature dendritic cells infected with ZIKV WT or mutants. (D) Quantification of viral genome copies in the infected immature dendritic cells. The relative viral genome copies to the WT virus was plotted as means ± SD (n=3).

## Discussion

In this study, we generated an infectious cDNA clone of ZIKV strain PRVABC59, and used a high-throughput fitness profiling approach to assess the impact of mutations at every amino acid position of E protein on viral fitness. We found that 9 resides showing high polymorphism in natural sequences could tolerate mutations, and we also evaluated the role of four conserved residues involved in E protein pH-induced conformation change. By comparing ZIKV replication fitness in mosquito and human cells, we discovered that mutations affecting N-linked glycosylation of E protein presented the most dramatic difference. While these mutations either only slightly or did not affect ZIKV infection of human cells, they significantly increased ZIKV growth in mosquito cells. In an attempt to explore the underlying mechanism, we discovered that ablation of N-linked glycosylation enhanced ZIKV entry into mosquito cells. Furthermore, N154 glycosylation was found to be important for ZIKV infection of mammalian cells through entry factor DC-SIGN, indicating functional conservation among flaviviruses.

### High-throughput fitness profiling of ZIKV E protein

Traditionally, mutagenesis at a specified position is the common approach for examining functional residues of a virus. The underlying principle of our high-throughput fitness profiling approach is to mutagenize the entire viral gene (or even genome) towards a single-nucleotide resolution, aiming to interrogate every position across the viral gene, passage the mutant library in a desired selection condition, and quantify the frequency change of each mutation to evaluate its phenotype. By using a mutant library of ZIKV E protein where every individual amino acid is substituted with other residues, this study represents the first application of a high-throughput approach to analyze the functional residues of an entire ZIKV protein. In the future, this high-resolution profiling approach could be applied to other proteins in order to systematically understand ZIKV infection and its related pathogenesis. Additionally, our library of high-dense ZIKV E mutants could be applied to examine roles of E protein in other aspects. In recent studies, mutant libraries of viral envelope proteins were used to characterize viral mutations escaping from monoclonal antibodies neutralization, and predicted antibody binding regions (Fulton et al., 2017; Lin et al., 2012; Michael B Doud, 2016). Through a similar strategy, our high-resolution profiling approach, facilitated by the library of ZIKV E mutants, could be applied to define protein epitope(s) for neutralizing antibody(s), including monoclonal antibodies and polyclonal antibodies in patient sera.

### N-linked glycosylation at position N154 of ZIKV E protein

Employing the high-throughput fitness profiling analysis, we are able to identify mutations that impair viral fitness in a specific cell background, which enables us to select informative mutants to dissect the multiple functions of a protein. In this study, by comparing ZIKV fitness in mosquito and human cells, we discovered that mutations affecting the N-linked glycosylation of E protein presented the most dramatic difference. In studies of other flaviviruses, N-linked glycosylation at position N153/154 of the E protein was shown to be important for viral growth in both mammalian and mosquito cells (Beasley et al., 2005; Hanna et al., 2005; Lee et al., 2010; Mondotte et al., 2007; Scherret et al., 2001). For example, ablation of N153 glycosylation of the E protein reduced DENV production over 10 folds in cultured cells, including mosquito cells (C6/36), mammalian epithelial cells (Vero) and fibroblast cells (BHK21) (Lee et al., 2010; Mondotte et al., 2007). Similarly, N154 glycosylation was required for the efficient growth of WNV in multiple cell lines, including C6/36, Vero, and DF-1 (Beasley et al., 2005; Moudy et al., 2009; Scherret et al., 2001). Surprisingly, N-linked glycosylation at position N154 of E protein was not necessary for ZIKV replication in either human epithelial cells (A549) or endothelial cells (hCMEC). In addition, ablation of N-linked glycosylation significantly enhanced ZIKV growth in C6/36 cells (Figs. 4, 6A & 6B). Further exploring the mechanism, we found that N-linked glycosylation of ZIKV E protein did not affect initial attachment of virions to the cell surface (Fig. 6C), but did notably impair viral entry into C6/36 cells (Fig. 6D). In a study of WNV using subviral particles (SVPs), SVPs lacking glycosylation of E protein were found to be modestly more efficient in entry on BHK-21 and QT6 cells, while this absence enhanced the infection of C6/36 cells. However, the latter experiments showed this increase of entry didn’t lead to more viral production of non-glycosylated mutants, probably due to deficiency in virion assembly and egress (Hanna et al., 2005).

The initial step in the life cycle of flaviviruses is attachment of the virion to host-cell entry factor(s), followed by entry into cells via endocytosis through which the virus particles are transported to endosomes. The acidic environment in the endosome lumen promotes rearrangement and conformational change of E protein, resulting in the fusion of the virion envelope with the endosomal membrane, followed by release of viral genome into cytoplasm (Heinz et al., 2004; van der Schaar et al., 2007; Zhang et al., 2004). Based on cryoEM structures of ZIKV, the glycans at N154 position of the E protein lie over the dimer interface in close proximity to the fusion loop of domain II (Kostyuchenko et al., 2016; Sirohi et al., 2016). Considering their substantial size, glycans might generate steric effects impairing ZIKV E protein conformational change in endosome and/or the membrane fusion process between virion and endosome of mosquito cells, which can explain why ablation of N-linked glycosylation enhanced ZIKV entry in C6/36 cells (Fig. 7D). However, we also found that ablation of N-linked glycosylation didn’t enhance ZIKV infection of human A549 and hCMEC cells, implying that it plays different role(s) in entry of mosquito and human cells. In future research, the mechanisms underlying the impairment of N-linked glycosylation of E protein on ZIKV entry in mosquito cells and the different roles of glycosylation among flaviviruses, need to be investigated.

A N154-D155-T156 sequence, required for N-linked glycosylation, exists in many but not all ZIKV strains (Faye et al., 2014; May and Relich, 2016). E protein variants defective in N-linked glycosylation, especially a Thr to Ile (T156I) change, were frequently observed among ZIKV stains. It is suggested that acquisition of the N154 glycosylation site is a recurrent event in the history of ZIKV (Faye et al., 2014). However, this hypothesis could be limited as some loss-of-glycosylation variants might be a consequence of passaging ZIKV strains before sequencing. In our study of ZIKV Puerto Rico strain, mutations of residue N154 did not obviously change viral growth in two human cells, significantly enhanced viral fitness in mosquito cells (Figs. 5 & 6), and impaired virus infection through DC-SIGN entry factor (Fig. 7). However, in ZIKV African isolates, substitution or deletion of N154 is rarely observed. In addition, we noticed that, when compared to the corresponding silent mutations, 2 out of 5 mutations (N154Y and N154K) of N154 did not obviously increase viral fitness in mosquito cells, while all six mutations of T156 did so significantly(Fig. 5A). All the above information suggests that residue N154 may also serve other function(s), independent of N-linked glycosylation, in ZIKV replication. In addition, they also validate the advantage of the high-resolution fitness profiling platform in unbiasedly analyzing functional residues.

Previously, it has been shown that DC-SIGN mediates flavivirus infection of human dendritic cells by binding E protein glycan(s) (Hamel et al., 2015; Tassaneetrithep et al., 2003), and dendritic cells were important targets for flavivirus infection in patients (Marovich et al., 2001; Wu et al., 2000). In addition, N-linked glycosylation of the E protein of WNV could possibly enhance its neuroinvasion (Beasley et al., 2005; Shirato et al., 2004). In our study, N-linked glycosylation of the E protein was shown to be important for ZIKV infection of immature dendritic cells and also 293T cells expressing DC-SIGN (Fig. 7). The above information suggests that glycosylation of ZIKV E protein may contribute to its virulence in patients and mouse models. However, all of the data in our study were collected from *in vitro* infection of human and mosquito cell lines, future detailed investigation with mouse models and mosquito vectors will be required to examine the potential connection between E protein N-linked glycosylation and ZIKV virulence, particularly neuroinvasion.

In summary, this is the first study to systematically map functional residues of the ZIKV E protein, and characterize the roles of its N-linked glycosylation at N154 position. The study validates the advantage of a high-resolution fitness profiling platform in unbiasedly analyzing functional residues. Future research will be required to elucidate the molecular mechanism of the viral entry impacted by N-linked glycosylation of ZIKV E protein, in cell culture and in animal models. Our library of ZIKV E mutants could be employed to examine roles of E protein in other conditions, for example mapping neutralizing antibody binding epitopes and interactions with entry factor(s). We also expect that the high-throughput fitness profiling approach will be applied to study other ZIKV viral proteins.

## Author Contributions

All experiments were performed by D.G. with the help of D.Z. and L.W.. D.G. and R.S. designed the project. D.G., T.Z and Y.D. analyzed the data. Y.S., T.J.C, D.C., G.Z., P.S. T.T.W. and V.A. provided reagents and advices. D.G. wrote the manuscript. T.J.C, L.W. and R.S. revised the manuscript.

## Acknowledgement

We thank Dr. Ralph Baric for sharing data and comments, members of the Ren Sun and Ting-Ting Wu labs for discussions. This work was supported by NIH grants DE023591 and CA177322.

## Material and methods

### Cells, viruses and plaque assays

*Aedes albopictus* C6/36 and *Aedes aegypti* CCL-125 cells were obtained from ATCC. C6/36 cells were maintained in DMEM medium supplemented with 10% FBS at 28 °C with 5% CO2; CCL-125 cells were maintained in MEM medium supplemented with 20% FBS. Vero and BHK21 cells were cultured in DMEM with 10% FBS. Human epithelial A549 cells were cultured in RPMI1640 with 10% FBS. Human cerebral microvascular endothelial cells (hCMEC/D3) were maintained in EndoGRO-MV complete culture media (Millipore). The parental ZIKV Puerto Rico strain PRVABC59 (GenBank number KU501215) was obtained from United States Centers for Disease Control and Prevention. Titers of infectious viral particles were measured by a standard plaque assay as descried previously (Qi et al., 2015). Briefly, 6X10^4^ Vero or A549 cells were seeded in 12-well plates one day before infection, then infected with virus for 1 h and overlaid with 1% methylcellulose (Sigma). Four days later, cells were fixed and stained with 0.2% crystal violet in 20% ethanol, and plaques were counted to determine the titer.

### Human dendric cells generation

Human PBMCs were isolated from healthy donors by density centrifugation over Ficoll-Paque Plus in UCLA virology core. Monocytes were isolate from PBMCs with CD14 magnetic beads (Miltenyi Biotec), and cultured at 10^6^ cells/ml in RPMI1640 medium supplemented with 10% human serum, 1000U/ml each of human GM-CSF and IL-4 for 7 days.

### Construction of ZIKV infectious cDNA clone

After receiving the virus from CDC, ZIKV was amplified in C6/36 cells, and total RNA of infected cells was extracted with Purelink RNA Mini Kit (Ambion). ZIKV cDNA covering the complete genome was synthesized using SuperScript III reverse transcriptase (Thermo Fisher) with random hexamer (Fragments 1 to 5) or ZIKV specific primer against the last 20 nt of ZIKV 3’-end (Fragment 6). Then, six ZIKV sub-genomic fragments covering the whole genome were amplified using KOD polymerase (Millipore) with primers listed in Table S1. Six fragments were assembled into EcoR I linearized pBR322 plasmid to generate pZ-PR plasmid using HiFi DNA Assembly Cloning Kit (NEB), and further transformed into DH10B E.coli. The whole sequence of pZ-PR plasmid was verified by Sanger DNA sequencing. Sequence of pZ-PR is available from NCBI GenBank under accession number KY583506 and also in supplemental Materials and methods.

### Reconstitution of ZIKV from pZ-PR plasmid

Recombinant ZIKV was generated from plasmid pZ-PR, containing the full-length cDNA of ZIKV Puerto Rico strain, using the previously describe method (Shan et al., 2016) with some modifications. In Brief, pZ-PR plasmid was amplified in DH10B E.coli and then purified using PureLink HiPure Plasmid Midiprep Kit (Thermo Fisher). To generate DNA template for RNA *in vitro* transcription, 100 μg of pZ-PR was linearized with BstBI, followed by end blunting with Mung Bean Nuclease (NEB). Then, the end-blunted template DNA was purified with phenol-chloroform, precipitated with ethanol, and resuspended in 20 μl RNase-free water. 5’-capped RNA was *in vitro* transcribed using mMESSAGE mMACHINE T7 kit (Ambion) with 1 μg template DNA and an additional 1 μl of 30 mM GTP solution. *In vitro* transcribed RNA was further purified with phenol-chloroform, precipitated with ethanol, resuspended in 100 μl RNase-free water, aliquoted, and stored in -80 °C freezer.

For cell transfection, 12 μg of RNA was mixed with 5X10^6^ BHK21 cells in 200 μl Electroporation Solution (Bioland), and electroporated in 4-mm cuvette with the GenePulser apparatus (Bio-Rad) at the setting of 240 V and 950 μF, pulsing once. After 10 min recovery at room temperature, transfected cells were resuspended in 10 ml warm culturing medium, incubated in cell incubator overnight, and then washed once and replenished with fresh medium. Three to four days post electroporation when ∼40% of cells showing CPE, supernatants were collected, span at 8000 g for 10 min at 4 °C to remove cellular debris, aliquoted and stored at -80 °C.

### Generation of the plasmid DNA library of ZIKV E protein mutations

To generate the mutant plasmid DNA library of ZIKV E protein, we designed an error-prone PCR strategy utilizing the pZ-PR plasmid as the template and the GeneMorph II Random Mutagenesis Kit (Agilent) to generate the point mutations during PCR amplification (Al-Mawsawi et al., 2014; Wu et al., 2015). The ZIKV E coding region was divided into three segments ranging from 516 to 545 bp (Fig. 2A) for the purpose of controlling mutational rate in error-prone PCR reaction. The mutated inserts were generated with 12-cycles amplification using the GeneMorph II Random Mutagenesis Kit, 1 μg pZ-PR plasmid and corresponding primers (Table S1). To generate full length fragments for cloning, the corresponding left and right fragments were amplified using high fidelity KOD DNA polymerase (Millipore) for 18 cycles. Each mutated insert and its corresponding left and right fragments were combined, and the resulting full length DNA inserts were generated by amplification with KOD polymerase for 20 cycles. Then, the full length inserts were digested with KpnI and EcoRI, while the vector pZ-PR plasmid was digested with KpnI and EcoRI followed by treatment with Shrimp Alkaline Phosphatase (NEB). Ligation was performed for each of the three sub libraries with T4 DNA ligase (NEB) with a vector: insert ratio of 1:5. Ligated products were purified with phenol-chloroform, precipitated with ethanol, resuspended in 10 μl sterilized water, and electroporated into DH10B E.coli competent cells. For each of the three sub libraries, ∼40,000 colonies were collected from LB plates, and directly subjected to plasmid DNA purification. Equal amounts of the three sub libraries were then mixed to generate the DNA library, and stored at -80 °C in aliquot.

### Preparation of virus library and passages

To reconstitute the virus library, 5’ capped RNA was *in vitro* transcribed from the plasmid DNA library, electroporated into BHK21 cells, and cell supernatant was collected at 4 days post transfection. Cell supernatant containing the library of virus mutants was span at 8000 g for 10 min at 4 °C to remove cellular debris, titrated and stored at -80 °C in aliquot. Virus library was further passaged in C6/36, A549 or hCMEC cells by infecting 40 million cells at a MOI of 0.05. Virus supernatants were collected at 4 days post infection, debris clarified, RNase A treated, and then subjected to RNA purification followed by deep sequencing analysis.

### Next-generation sequencing of virus mutations and data analysis

Viral supernatants were treated with RNase A, then virion RNA was extracted using QIAamp Viral RNA Mini Kit (Qiagen), and reverse transcribed to cDNA using Superscript III Reverse Transcriptase (Thermo Fisher). The plasmid DNA library or cDNA from the viral libraries (virus or passaged libraries) were used as templates to generate thirteen158-bp deep sequencing amplicons with primers listed in Table S1. The resulting PCR amplicons were 3’-end dA-tailed, and ligated to Y-shape sequencing adapters using T4 DNA ligase (NEB). Y-shape adapters were generated by annealing two oligos: 5’-ACA CTC TTT CCC TAC ACG ACG CTC TTC CGA TCT NNN NNN T-3’ and 5’-/5Phos/NNN NNN AGA TCG GAA GAG CGG TTC AGC AGG AAT GCC GAG-3’. The six “N” stands for the multiplex ID for distinguishing different samples. The adapter-ligated products were further PCR amplified for 18 cycles using KOD polymerase and UniFcell primers (Table S1). The final PCR products were gel purified, and pooled for deep sequencing on an Illumina Hiseq 3000 platform with 150 bp paired-end reads. Raw sequencing reads were demultiplexed by using the six-nucleotide barcodes. Sequencing error was corrected by filtering unmatched forward and reverse reads. Each mutation was called by comparing individual reads to the WT pZ-PR reference sequence. The relative frequency and relative fitness score (RF score) for a given mutation was calculated as follows: For a mutation *i* in deep sequencing amplicon *n* of sample *t* (where *t* could be DNA plasmid, or virus libraries):

Relative frequency_*i,n,t*_ = Read count_*i,n,t*_ / Coverage_*n,t*_, where Read count_*i,n,t*_ represented the number of read in amplicon *n* of sample *t* that carried mutation *i* and Coverage_*n,t*_ represented the total read count (including WT and all mutations) of the amplicon *n* of sample *t*. RF score = Relative frequency_*i,n,Virus*_ / Relative frequency_*i,n,DNA*_, where Relative frequency_*i,n,Virus*_ represented relative frequency of mutation *i* in viral library (virus or passaged library) and Relative frequency_*i,n,DNA*_ represented relative frequency of mutation *i* in DNA plasmid library.

All analysis was performed with customized python scripts, which are available upon request. Raw sequencing data is available from NIH Short Read Archive under accession number PRJNA373927.

### Polymorphism calculation

Polymorphism score of each residue was calculated using a formula previously reported (Crooks et al., 2004). S = -100 * Sum (Pi * logPi) where Pi is the frequency of mutations at position i. The score is the normalized entropy of the observed mutation distribution.

### Construction of individual ZIKV mutants

Single nucleotide mutations were introduced into pZ-PR plasmid individually, using PCR-based site-directed mutagenesis strategy with primers listed in Table S1. To generate mutant virus, BHK21 cells were electroporated with *in vitro* transcribed RNA, and viral supernatant were collected at 4 days post transfection. Mutant viruses were further amplified in C6/36 cells, then supernatants were collected, clarified of debris and stored at -80 °C in aliquot. Virion RNA was also extracted, and reverse transcribed to cDNA. E protein coding sequence of all mutants were PCR amplified individually, followed by Sanger DNA sequencing to verify the sequence.

### Reverse transcription and real-time PCR

Quantitative RT-PCR was performed as previously described (Gong et al., 2016). Briefly, total RNA was extracted from infected cells with Purelink RNA Mini Kit (Ambion) or from RNAase A treated cell supernatant with QIAamp Viral RNA Mini Kit (Qiagen), reverse transcribed using Superscript III Reverse Transcriptase (Thermo Fisher) and random hexamer, and then subjected to real-time PCR analysis using primers: 5’-TTG TGG AAG GTA TGT CAG GTG-3’ and 5’-ATC TTA CCT CCG CCA TGT TG-3’.

### Western blotting

ZIKV infected C6/36 or A549 cells were lysed in denaturing buffer, mock treated or treated with PNGase F(NEB) to remove N-linked glycan from proteins. Then proteins were diluted 5 times in SDS-PAGE sample buffer, heated at 95 LJC, resolved by SDS-PAGE gel electrophoresis, and transferred onto PVDF membrane. ZIKV E protein was detected with the monoclonal antibody D1-4G2-4-15 (Absolute Antibody). HRP-conjugated secondary antibodies were used and detection was performed with SuperSignal West Femto Maximum Sensitivity Substrate (Thermo Fisher).

### Indirect immunofluorescence assay

Expression and localization of ZIKV E proteins was determined by indirect immunofluorescence assay. Cells were infected with ZIKV WT or mutants, fixed in 2% paraformaldehyde, permeabilized with 0.1% Triton-X100, and then blocked with 3% BSA plus 10% FBS. ZIKV E proteins were detected with mouse monoclonal antibody clone D1-4G2-4-15 (Absolute Antibody), followed by Alexa Fluor 488-conjugated anti-mouse IgG antibody (Life Sciences). DAPI was used for the staining of DNA. Samples were examined using a Nikon microscope using 63X/1.40-0.60 oil lens.

### Statistical analysis

All numerical data were calculated and plotted with mean+/-SD. Results were analyzed by unpaired Student’s *t* test. Differences were considered statistically significant when p<0.05 (*) or p<0.01 (**).

